# Inhibitory and excitatory mechanisms in the human cingulate-cortex support reinforcement learning

**DOI:** 10.1101/318659

**Authors:** Vered Bezalel, Rony Paz, Assaf Tal

## Abstract

The dorsal anterior cingulate cortex (dACC) is crucial for motivation, reward- and error-guided decision-making, yet its excitatory and inhibitory mechanisms remain poorly explored in humans. In particular, the balance between excitation and inhibition (E/I), demonstrated to play a role in animal studies, is difficult to measure in behaving humans. Here, we used magnetic-resonance-spectroscopy (^1^H-MRS) to examine these mechanisms during reinforcement learning with three different conditions: high cognitive load (uncertainty); probabilistic discrimination learning; and a control null-condition. Subjects learned to prefer the gain option in the discrimination phase and had no preference in the other conditions. We found increased GABA levels during the uncertainty condition, suggesting recruitment of inhibitory systems during high cognitive load when trying to learn. Further, higher GABA levels during the null (baseline) condition correlated with improved discrimination learning. Finally, excitatory and inhibitory levels were correlated during high cognitive load. The result suggests that availability of dACC inhibitory resources enables successful learning. Our approach establishes a novel way to examine the contribution of the balance between excitation and inhibition to learning and motivation in behaving humans.

## Introduction

Studies in animals have highlighted the importance of excitation and inhibition for reinforcement learning[1-3]. The balance between them (E/I balance) is critical and maintained under most conditions, yet the exact ratio is highly dynamic[4, 5], and variations support information processing and learning[4, 6]. Although it is much harder to assess excitation and inhibition in humans, the main contributors: Glutamate and GABA, can be accurately and reliably measured through Proton Magnetic Resonance Spectroscopy (^1^H-MRS) [7-9]. Indeed, MRS-observed neurotransmitter levels reflect task-related activity with studies demonstrating sensitivity to baseline [10-15] and rapidly-modulating levels [16-18]. Most studies measured concentrations during rest and correlated it with later/previous behavior[13-15, 19-21], yet some even measured during behavior[17, 18, 22]. A few studies have correlated baseline (rest) levels with subsequent/prior learning metrics[11, 12, 23], and one has even examined changes during motor learning[16]. However, none have quantified neurotransmitter modulations during active reinforcement-learning.

Error-based learning involves the dorsal-anterior-cingulate-cortex (dACC), which mediates motivation, cognition and action[24-28]. The dACC plays a crucial role in reward-guided decision making, as it promotes action-outcome associations and monitors goal-directed behaviors [27, 29]. In particular, dACC activation is modulated by requirements for cognitive control [30-32], and is involved in monitoring choice outcome in uncertain environments [33-35], as well as biases decisions that require high mental effort [36-38]. However, the contribution of excitation-inhibition mechanisms to these functions in the human ACC remains poorly understood.

We used ^1^H-MRS during reinforcement learning in humans, and measured modulations in dACC levels of GABA and Glx while participants engaged in a learning task that compared three factors: full uncertainty (high-cognitive-load); probabilistic discrimination learning, and an active Null condition. We hypothesized that levels of neurotransmitters during the high-cognitive-load, and potentially during active-Null conditions, would predict and enable successful discrimination learning.

## Results

A two-alternative forced choice task was conducted using a block design paradigm while subjects (n=31) laid in an MRI scanner (Fig.1A). During each trial, subjects heard two pure-frequency tones, each to a different ear, in pseudorandom order. Following a visual go-signal comprised of two arrows, the subjects chose a tone by pressing right or left button. The outcome was one of three: a monetary gain (+2), a loss (−2), or neutral feedback (0). There were three experimental blocks: (1) the Null condition where the subjects received neutral (0) outcome, independent of their choice; (2) the uncertainty condition, 50/50 probability for loss/gain, independent of the participant’s choice; and (3) the probabilistic discrimination condition, in which one of the tones was paired with a monetary gain in 80% of the trials and monetary loss in 20%, and vice versa for the other tone. All the subjects underwent a rest scan before the experimental blocks. Additionally some subjects (n= 18) also underwent a rest scan after the experimental blocks.

**Figure 1.**
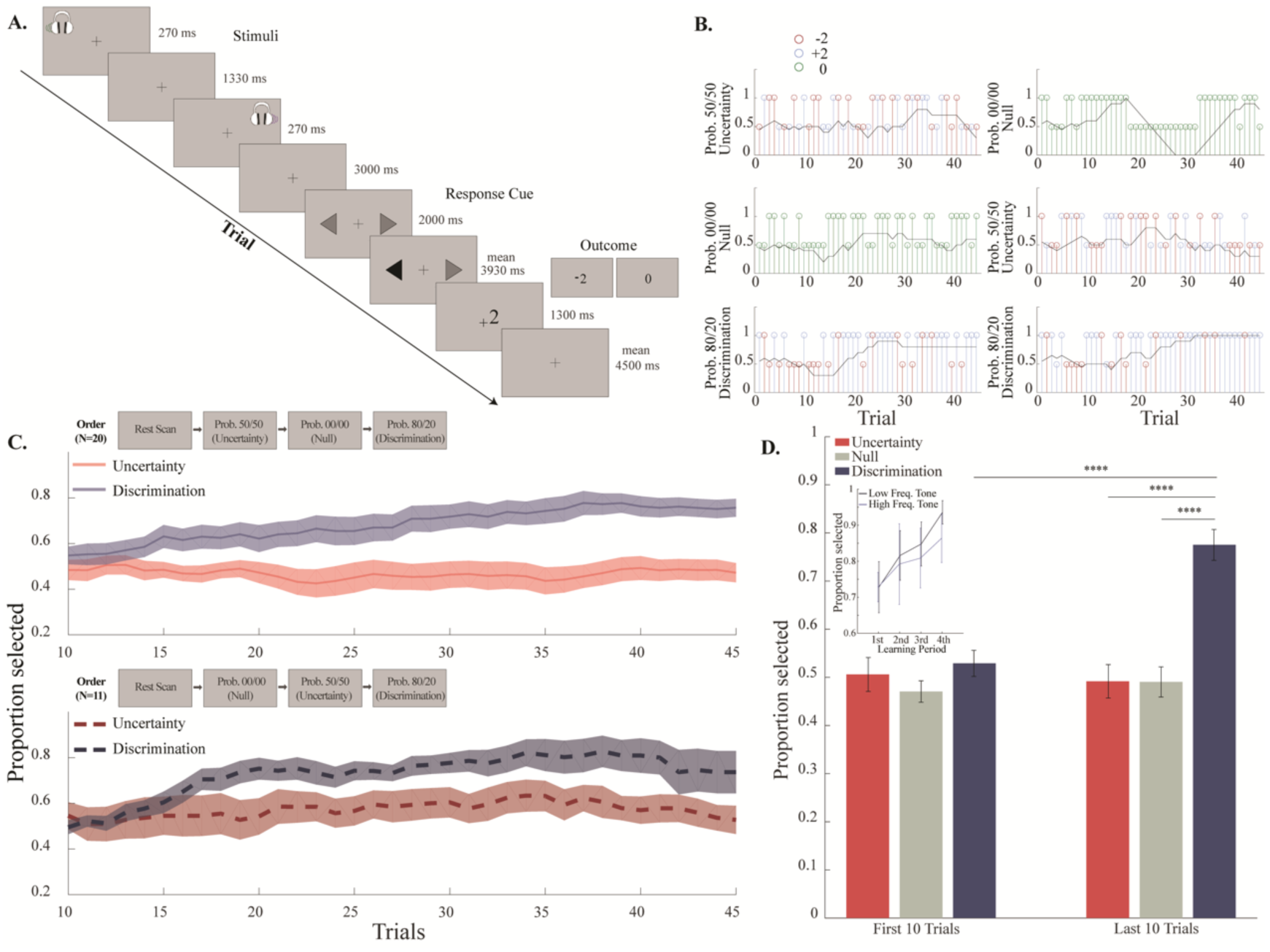
Experimental design and behavioral results. **A.** In each trial, subjects were exposed to two pure tones that were played out in succession to different ears. The response cue, presented by two gray opposing arrows, instructed to choose between the tones (left or right button press). Following selection, the arrow corresponding to the chosen laterality was blackened, and the outcome screen indicated monetary gain, loss or neutral outcome (+2,-2 or 0). **B.** Behavior of single two subjects (rows), in the order of exposure to each of the 3 conditions (rows, top to bottom). The scanning session consisted of three scanning blocks, each attributed to one behavioral condition: 50/50 probability to lose or gain 2₪ (“uncertainty condition”), 00/00 probability with consistent 0₪ reward (“Null condition”), and 80/20 probability to lose or gain 2₪ (“discrimination condition”). Blue, red and green represent gain, loss and zero reward respectively. Taller stems represent selection of one tone, while shorter stems represent selection of the other tone (1 and 0.5 in the y axis respectively). The black line is the probability to choose the tone designated as the gain-tone in the 80/20 condition, averaged over a 10-trial moving window. **C.** Tone selection probability during the uncertainty and discrimination conditions, presented separately for the two ordering of conditions (mean +/- shaded SEM). Top: average across subjects that were exposed to the first ordering (n=20): MRS rest scan, followed by the uncertainty, Null, and discrimination conditions. Bottom: average across subjects that were exposed to the second ordering of conditions (n=11): MRS rest scan followed by the Null, uncertainty, and discrimination conditions. The graphs demonstrate a gradual increase in the high-gain-tone (‘better-option’) selection probability during the discrimination condition, irrespective of the ordering, and no selection preference during the uncertainty condition (see text for statistics). **D.** High gain tone selection probability averaged across the first and last 10 trials in each condition (mean +/- SEM). Participants chose the high gain tone more often in the last part of the discrimination condition compared to its first part, as well as to the last parts of the two other conditions (\*\*\*\**p*<*0.0001*, post-hoc Tukey-Kramer). Inset presents pretest results (n=10) showing that learning was similar regardless of whether a high frequency (light blue) or low frequency tone served as the better-option (Wilcoxon *p*>*0.05* in each one of the four points comparisons).

### Learning occurs under probabilistic discrimination, but not under uncertainty or Null

Over the course of the discrimination block, subjects gradually increased their preference to the tone that was associated with higher gain (‘better-option’), such that by the end of this block, the mean selection probability exceeded 70% (Fig.1B,C). In comparison, no preference to either one of the options was observed during the other two conditions, in which the subjects presented a stable mean probability of 0.5 ± 0.04 (uncertainty condition) and 0.48± 0.03 (null condition) to select each option (Fig.1B,C). This was the case for both groups of subjects, when the uncertainty condition was first (Fig.1B-left, Fig.1C-top, n=20), and when the Null condition was first (Fig.1B-right, Fig.1C-bottom, n=11).

The preference to the better-option in the discrimination condition was also reflected by the averaged selection probability during two epochs, namely, the first and last 10 trials of the block (Fig. 1D). A difference in selection probability between conditions was observed (p<0.00001, F (2, 60) = 18.3, two way repeated measures ANOVA; mean selection probability of the first and second epochs: 0.5, 0.48 and 0.65, in uncertainty, null and discrimination conditions, respectively), arising from a significant preference to the better-option in the discrimination condition when compared to the uncertainty and null conditions (p<0.001 and p<0.00001 respectively, post hoc Tukey-Kramer). Additionally, we also observed a difference between the first and second epochs (p<0.001, F (1, 30) = 19.2, two way repeated measures ANOVA) that originated from a preference to the better-option in the second epoch (p<0.001, post hoc Tukey-Kramer). An interaction between these effects was observed (p<0.0001, F (2, 60) = 12.2, two way repeated measures ANOVA), deriving from the increased preference to the higher gain tone in the second epoch of the discrimination condition compared to its first epoch (p<0.00001, post hoc Tukey-Kramer). Examination of the performance in the second epoch revealed a preference to the high-gain-tone in the discrimination condition compared to the null and uncertainty conditions (both p<0.00001 post hoc Tukey-Kramer). However, there was no difference in the average selection probability between the conditions in the first epochs of the blocks (p>0.05, post hoc Tukey-Kramer).

We conclude that subjects learned the task in the probabilistic discrimination condition, behaved at chance level in the Null condition (no outcomes), and also behaved at chance level under uncertainty (when rewards and punishments occur, but there is no reliable information to learn, leading to a high-cognitive-load).

### Measuring neurotransmitter concentrations in the dACC using ^1^H-MRS

The MRS technique requires long scanning sessions in order to achieve good signal to noise ratio. Therefore, we defined the region-of-interest (the dACC) a-priori based on anatomical maps (Fig.2A). Please see methods for full description of how we measure and quantify GABA and Glx concentrations (Fig.2B). In addition, we conducted two tests to assess whether results might be confounded by the order of experimental conditions and/or the stability of metabolite levels over the long scanning time.

**Figure 2.**
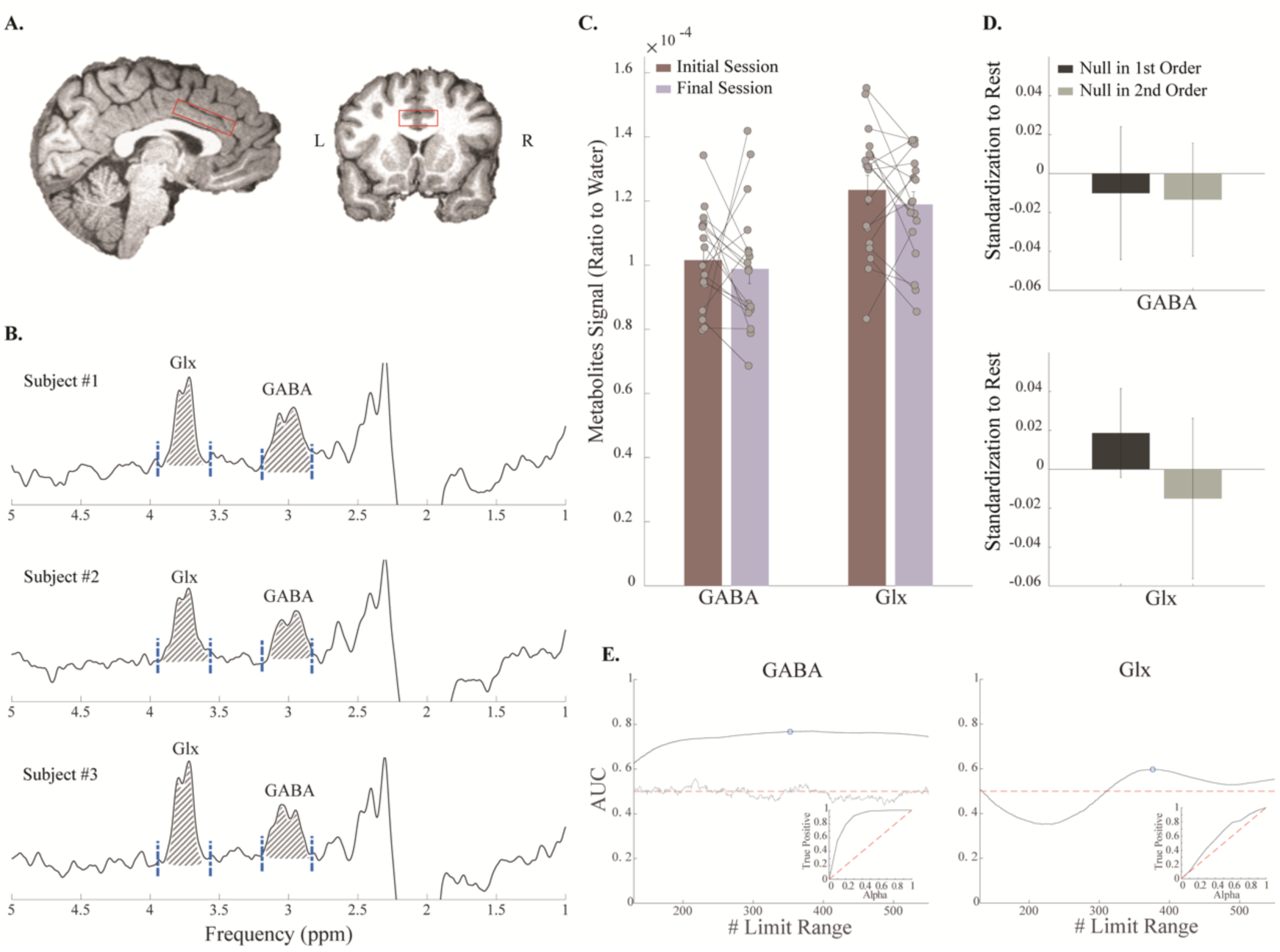
MRS measurements of GABA and Glx concentrations are stable. **A.** A 40×25×10 mm^3^ voxel was placed on the midline of the dACC, chosen anatomically, presented in sagittal (left) and coronal (right) T1-weighted anatomical MPRAGE scans. **B.** Sample spectra from three different subjects (144 averages, 9:57 minutes), displaying the peaks of GABA and Glx at 3.01 and 3.75 ppm respectively. Blue dashed lines represent the integration limits that were used to quantify each metabolite. The integrated area under the curve, used for quantification, is illustrated by diagonal lines. **C.** Eighteen subjects went through an additional final session of rest scan after completion of the task. Comparison between metabolites levels in the initial and in the final rest scans, revealed no difference over time in GABA/Water and Glx/Water levels (*p*>*0.05*, paired sample t-test). Levels of metabolites in single subjects are represented by connected dots (mean +/- SEM). **D.** Order of experimental conditions did not affect metabolite levels in the Null condition. Compared between the first (n=20) and the second (n=11) orderings for both GABA and Glx standardized to rest (Wilcoxon *p*>*0.05*). (mean +/-SEM). **E**. Chosen integration limits for each metabolite were validated to be robust by an optimization process. We varied the limits and used a ROC curve analysis (insets) to choose the best limits. Shown are examples of the area-under-curve (AUC) for each pair of tested integration limits, with the best integration limits for each metabolite circled in blue. The grey line is based on simulated random noises as control. The results are highly similar to the limits chosen traditionally (usually based on visual considerations alone).

First, we assessed putative temporal metabolic drift by examining metabolites levels during the rest scan, which is the only scan type that was not influenced by task manipulation but could be affected by time. Metabolites levels were compared in subjects that performed the rest scan block twice: at the beginning and at the end of the experiment (Fig.2C, n=18). There was no difference between the levels of GABA/Water (t (17) = 0.46, p>0.05, paired sample t-test) or Glx/Water (t (17) = 1.06, p>0.05, paired sample t-test). We conclude that the time did not influence metabolite levels and is not a major contributor to the results.

To further validate that the order of conditions was not a factor, we had two groups of subjects, one performed the uncertainty before the Null condition (n=20), and the other performed the Null condition before the uncertainty (n=11), and compared levels of GABA and Glx during the Null condition in the two experimental orders. Because we already established that metabolite levels during rest are stable from start to end, we standardized GABA and Glx levels during the Null condition to the initial rest level (Eq. (1)). We found no difference (Fig.2D) between the signals of GABA (Wilcoxon p>0.05, Z=0.14) or Glx (Wilcoxon p>0.05, Z=0.56).

### Increased GABA during uncertainty condition only

Differences between the conditions of the task were found when examining GABA/Water and Glx/GABA levels (Fig. 3A; p<0.05 with F (2, 60) = 3.6 and F (2, 60) = 4.1 respectively, one way repeated measures ANOVA). These differences originated from elevated GABA/Water and reduced Glx/GABA levels during the uncertainty condition compared to the discrimination condition (p<0.05, post hoc Tukey-Kramer). Glx/GABA displayed a marginal non-significant trend for the difference between the Null condition to the uncertainty condition (p=0.052, post hoc Tukey-Kramer), but not for GABA/Water levels (p>0.1, post hoc Tukey-Kramer). No differences were found for GABA/water and Glx/GABA between the Null and discrimination conditions (p>0.1, post hoc Tukey-Kramer). Finally, Glx/Water did not exhibit any differences between the conditions (p>0.1, F (2, 60) = 0.5, one way repeated measures ANOVA).

**Figure 3.**
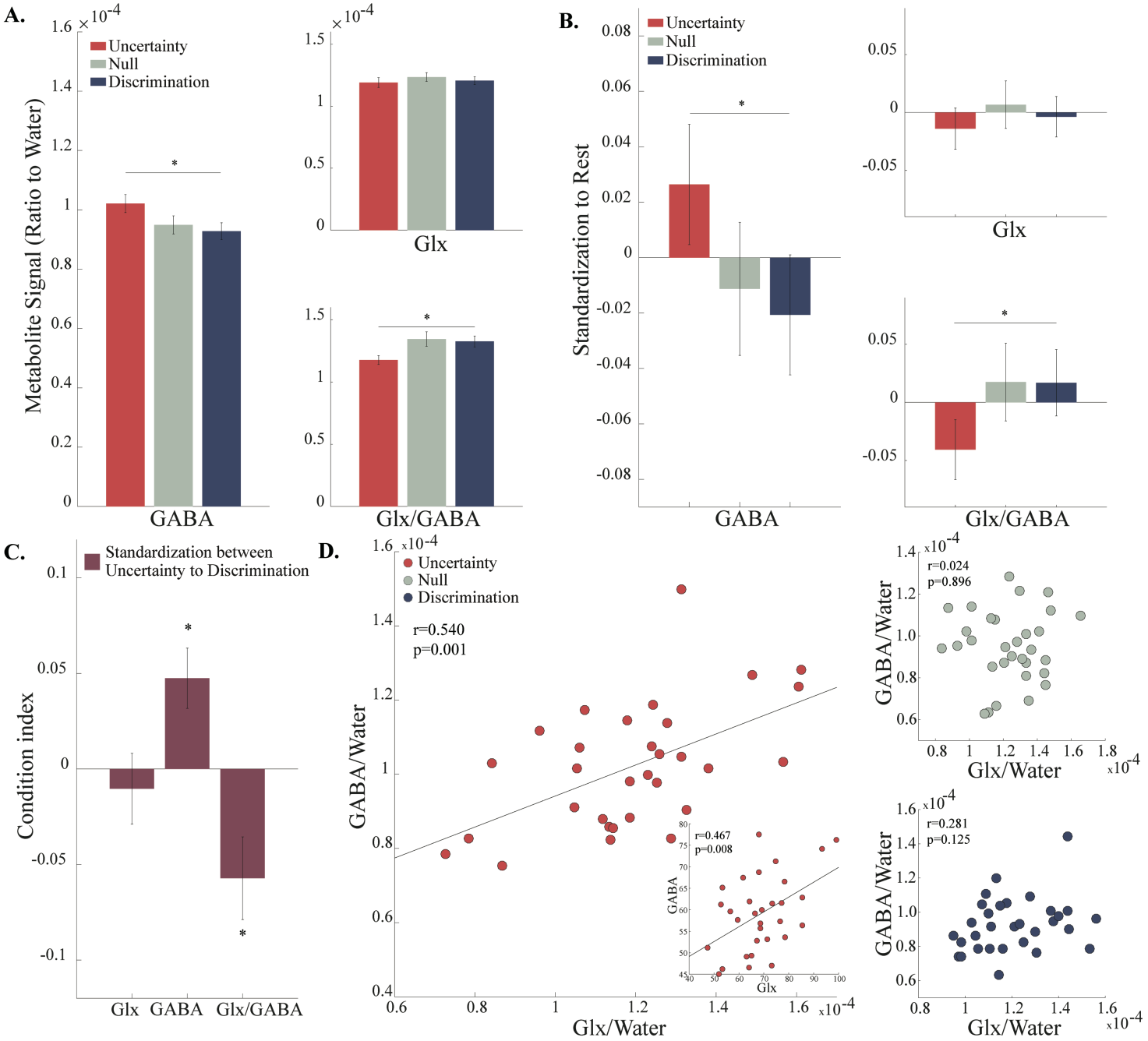
Higher GABA levels observed during the uncertainty condition. **A,B**, Metabolites levels quantified as ratio to water (A) or standardized to rest (B) reveal a consistent result: higher GABA levels and lower Glx/GABA levels in the uncertainty condition, yet similar lower levels during the Null and discrimination conditions (*p*<*0.05*, repeated measures one way ANOVA; \**p*<*0.05*, post hoc Tukey-Kramer). See text for detailed statistics. **C**. Standardization between pairs of task conditions reveal significant differences in GABA and Glx/GABA levels when comparing between the uncertainty and the discrimination conditions (\**p*<*0.015*, one sample t-tests with Bonferroni adjusted alpha; error bars represent the SEM). **D**. A positive correlation between Glx/Water and GABA/Water levels during the uncertainty condition only (*r*=*0.540*, *p*=*0.001*). Inset shows the same without normalization by water signal (*r*=*0.467*, *p*=*0.008*).

For robustness, we repeated these comparisons after standardization to rest levels, for Glx, GABA and Glx/GABA (Eq. (11)). This yielded similar results to the ones obtained using the raw data, mainly presented a significant difference between the uncertainty to the discrimination condition in GABA and in Glx/GABA levels. When standardized to rest, GABA and Glx/GABA levels were found to vary significantly between the uncertainty and discrimination conditions (Fig. 3B; p<0.05 with F (2, 60) = 3.5 and F (2, 60) = 3.8 respectively, repeated measures one way ANOVA; p<0.05, post hoc Tukey-Kramer). A non-significant trend was exhibited by Glx/GABA when comparing the uncertainty and Null conditions (p=0.086, post hoc Tukey-Kramer), but no differences were found when examining GABA (p>0.05, post hoc Tukey-Kramer). Additionally, no difference was found between the Null condition to the discrimination condition in GABA level or in Glx/GABA (p>0.05, post hoc Tukey-Kramer). No difference was found between the conditions in the Glx levels (p>0.05, F (2, 60) = 0.6, one way repeated measures ANOVA).

Furthermore, we standardize directly between conditions (see methods), and used one sample t-test with Bonferroni adjusted alpha level of 0.015 (0.05/3). The comparison between the uncertainty to the discrimination conditions (Fig. 3C) yielded a significant standardized GABA and Glx/GABA scores (t (30) = 3.0, p=0.005, and t (30) =-2.64, p=0.013, respectively; one sample t-test). As expected, Glx did not change significantly in this comparison (t (30) = −0.56, p>0.015, one sample t-test). The comparisons between the null to the uncertainty condition and between the null to the discrimination condition were non-significant in all measurements (p>0.015, one sample t-test, data not shown).

It was tempting to hope that Glx/GABA results have an additional component to the GABA findings alone, thereby providing evidence to a specific E/I balance mechanism. To assess whether the significant differences in Glx/GABA between the uncertainty and the discrimination conditions could be explained by GABA, Glx or their combined effect, we implemented repeated measures ANOVA on GABA/water with and without Glx/water as a varying covariate. The resulting effect size increased from η^2^=0.24 without the covariate to η^2^=0.26 with the covariate. Subsequently, the square of each effect size was modeled as an index of correlation between the learning conditions to the GABA measurement, and a test for the difference between correlations was performed on the Fisher-transformed correlations. The test revealed no-difference between correlations, meaning Glx measurement was not found to add any significant information (Z=0.07, p>0.05).

We conclude that the major changes between conditions occur due to changes in GABA concentrations mainly.

### GABA and Glx levels are correlated during uncertainty condition

Although Glx was not found to significantly contribute to the differences observed between the conditions in Glx/GABA, and these were driven mainly by GABA changes – we aimed to further analyze the relationships between Glx and GABA. To do so, we calculated Pearson’s correlation coefficient between Glx/Water and GABA/Water in each of the conditions (Fig. 3D). A positive correlation was found during the uncertainty condition (r=0.540, p=0.001), but not during the Null (r= 0.024, p>0.05) or discrimination (r=0.281, p>0.05) conditions. This persisted when examining the correlation between GABA and Glx without the contribution of water (Fig. 3D inset r=0.467, p=0.008), and again there were no significant correlations in the other two conditions (p>0.05 in both, r=-0.12 for Null condition; r=0.26 for discrimination condition). A regression of GABA on Glx levels showed that although they increased in correlation, the E/I balance does change (*β* = 0.4182, significantly different than 1, p<0.01).

We propose that changes in Glx, although too small to show significant variations across conditions, still maintain a balance with the GABA signal when it varies extensively and significantly i.e. when GABA levels are especially high as in the uncertainty condition.

### GABA levels in the Null condition predict learning performance

We tested whether GABA and Glx levels during each experimental condition were directly related to subjects’ behavioral performance during the discrimination condition, as a proxy for their learning ability (Fig.4A). To do so, we calculated Pearson’s correlation coefficients between the behavioral measures during the discrimination condition to GABA/water and Glx/water levels during the different conditions.

**Figure 4.**
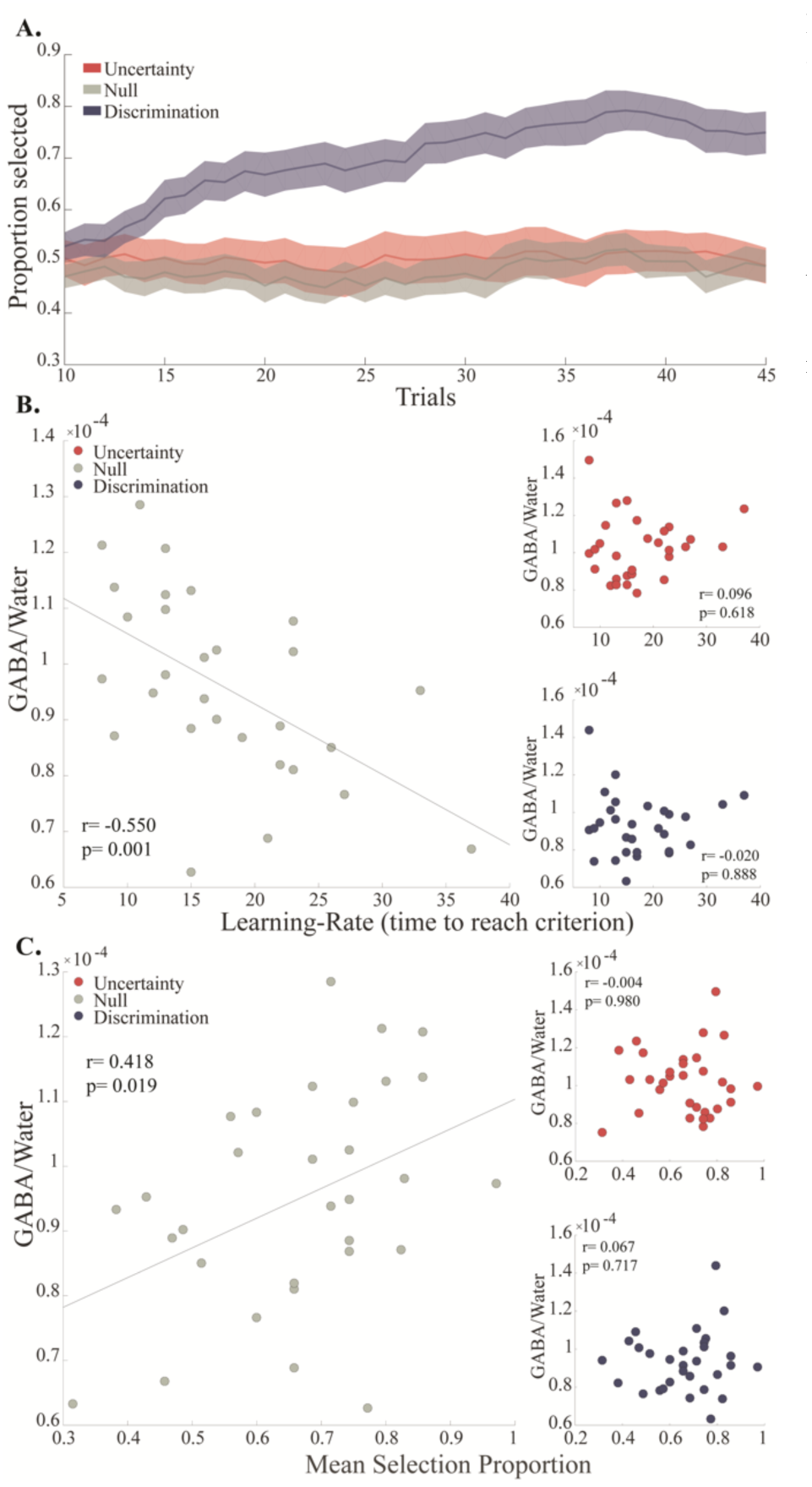
GABA levels in the Null condition predict behavioral performance during discrimination learning condition. **A.** Behavioral performance across all conditions. Gradual increase in selection probability of the better-option (high-gain tone) is observed during the discrimination condition. Averaged across all subjects (n=31) in 10-trial running window (mean +/-SEM). **B.** Negative correlation between the learning-rate, quantified as the trial # when a 0.7 criterion is reached during the discrimination condition, and GABA/water levels during the Null condition (*r*=-*0. 550*, *p*=*0.001*). **C.** Positive correlation between mean selection of the better-option during the discrimination condition and GABA/water levels during the Null condition (*r*=*0.418*, *p*=*0.019*).

We used two behavioral measures. First, we approximated the learning-rate as the trial when a subject reached performance criterion of 0.7, which is similar to the overall mean selection probability (0.69). There was a negative correlation (Fig.4B) between the learning-rate and GABA/water levels during the Null condition (r=-0.550, p=0.001), but not during the discrimination (r=-0.020, p=0.888) or uncertainty (r=0.096, p=0.618) conditions. Glx/water levels in the different experimental conditions did not reveal significant correlations with learning rate (p>0.05 in all conditions, r=-0.07 for Null condition; r<0.001 for uncertainty condition; r=-0.1 for discrimination condition).

Second, we assessed behavioral performance by the proportion of selection of the better-option. We found a positive correlation (Fig.4C) between the proportion and GABA/water levels during the Null condition (Fig.4C; r=0.418, p=0.019); but not during the other two conditions (r= −0.004, p=0.980 for uncertainty; r=0.067, p=0.717 for discrimination). As before, the selection probability was uncorrelated with Glx/water levels in any of the conditions (p>0.05 in all conditions; r=0.03, 0.09 and 0.15 for the null, uncertainty and discrimination conditions, respectively).

We conclude that baseline levels of available GABA, i.e. in a Null condition, predict better performance in a subsequent discrimination learning task.

## Discussion

The paradigm presented here simultaneously examined excitatory and inhibitory modifications in the dACC during learning. Behaviorally, the participants performed the task as expected, and learned to prefer the option that predicted higher gain in the discrimination condition, while lacked any preference in the two other conditions. These behavioral differences were expressed as increased GABA and decreased Glx/GABA levels during the uncertainty condition – when rewards and punishments are abundant, but there is no information to learn from. Although Glx was not significantly altered across conditions and did not seem to contribute beyond GABA to the difference in Glx/GABA across the conditions, further analysis revealed that Glx and GABA were positively correlated in the uncertainty condition only. This further supports the notion that when GABA is significantly increased, Glx levels do try to maintain a balance. Finally, examination of the relationship between GABA levels to the behavioral measures revealed that higher GABA levels in the Null condition predicted better learning, both when measured as time to reach criterion (learning rate), and when measured as final performance.

As previously shown, E/I balance is essential for cortical network stability [4, 5, 39, 40]. The precision of the balance depends on the degree of correlation between excitation and inhibition, ranging from a global balance in the absence of correlated inputs to a detailed balance for strong correlations [41]. The tight correlation, observed during the uncertainty condition, suggests a more homeostatic activity, in which specific functional patterns are amplified [42] and promote the occurrence of a distinctive functional activity. Moreover, we observed elevated GABA levels that can be interpreted as increase in inhibition which occurred in situations that are known to modulate the activity of the dACC, and consist of uncertain rewards [33-35, 43], high mental effort [36-38, 44], and requirements for cognitive control [30-32]. Additionally, a line of evidence linked the inhibitory system with achievement of efficient performance during high cognitive load [45, 46]. Therefore, presumably, during a need to make a decision in highly demanding cognitive load, specific functional patterns are amplified in the dACC and an increased inhibition occurs. This inhibition is required in order to achieve efficient and better performance. Similarly, individual differences in inhibition levels in the dACC might be linked to each individual experienced cognitive load, and reflect mental effort, uncertainty or a need for cognitive control.

In a neutral situation, such as the Null condition, the influences of external cognitive aspects of learning like incentives (rewards/punishments) are likely eliminated, exposing the underlying intrinsic motivational or attention-oriented processes, or simply the availability of the relevant neurotransmitter. Therefore, the correlation between inhibition in the dACC during neutral situation and later performance when learning is possible, might reflect the connection between the cognitive load and the motivational processes that take place during learning [47-50], which in their turn, lead to a better performance [51]. The fact that we did not identify any changes during learning itself (the discrimination condition) remain to be explored further. A possible interpretation is that learning occurred too fast for the long scan process required for MRS to detect changes beyond the signal-to-noise ratio; yet another possibility is that once uncertainty is reduced (because the subjects unveil a pattern to learn), less and less inhibitory mechanisms are required to maintain high cognitive load, attention, or uncertainty. A final possibility we cannot rule-out, is that the number of errors as well as punishments diminishes quickly, and the dACC signal diminishes with it. If either of the above is indeed the case, a design with much slower learning rate might allow detection of GABA levels in the dACC that are inversely-correlated with the learning performance.

Overall, the effects demonstrated here emphasize the importance of the dACC in processes of decision making and expand the understanding regarding its involvement in high cognitive demanding situations, and how it can later enable better learning. Importantly, it paves the way to further use of magnetic-resonance-spectroscopy (MRS) in identifying specific neural mechanisms and the use of neurotransmitters in brain networks during active behavior, and next steps should examine further regions, either by a-priori selection as done here, or by development of better sequences that allow simultaneous measurements from several regions. Combined with the paradigm we used here, it unveils the contribution of excitatory and inhibitory modifications during active learning in humans.

## Acknowledgments

We thank Dr. Edna Furman-Haran, Nachum Stern and Fanny Attar for assisting with the MRI procedures. This work was supported by ISF #26613 and ERC-2016-CoG #724910 grants to R. Paz, and the Minerva Foundation for A. Tal. A. Tal also acknowledges the support of the Monroy-Marks Career Development Fund and the historic generosity of the Harold Perlman Family.

## Methods

### Participants

All studies were performed in accordance to the procedures approved by the Internal Review Board of Haemek Medical Center (Afula, Israel). Thirty-seven right-handed healthy subjects (age range 22–39 years; median age 26 years; 21 females) participated in the experiment after written informed consent was obtained. Three subjects were excluded from the experiment due distorted signals, resulting from magnetic field drift, improper shimming or motion. Three additional subjects were excluded due to recurrent GABA and Glx outlier levels (more than 2.5 standard deviations).

### Behavioral Paradigm

A two-alternative forced choice task was conducted using a block design paradigm (Fig. 1A), while subjects laid in an MRI scanner. During each trial, subjects heard two short (270 ms) pure-frequency tones which were played out in succession and in a random order, each to a different ear. Subsequently, a visual response cue appeared, comprised of two arrows, corresponding to the two sides the tones were played to. The subjects chose their preferred tone by selection of the right or left remote buttons of a response box, corresponding to the side their preferred tone was played to. Following the selection, the chosen arrow was marked in black and subsequently an outcome screen appeared, presenting the response for the last selection, which was one of three possible outcomes: a monetary gain (+2), loss (−2) or neutral feedback (0). The paradigm consisted of three experimental conditions, each implemented within a separate block composed of 45 trials and lasting 12 minutes: (1.) the null condition with a consistent 0 reward (00/00 probability of loss/gain); (2.) the uncertainty condition (50/50 loss/gain probability, independent of the participant’s choice); and (3.) the discrimination condition (80/20 loss/gain probability), in which one of the tones was paired with a monetary gain in 80% of the trials it was chosen and monetary loss in 20% of the trials it was chosen, and vice versa for the other tone. Stimuli were generated by MATLAB (R2015b, The MathWorks, Natick, USA) using the Psychophysics Toolbox extension[52]. Each of the conditions was assigned one of three pairs of tone frequencies, mixed between subjects in pairing order. The tones’ frequencies were kept at a 4:7 ratio to facilitate their distinction. A fourth frequency pair was assigned to a short initial training session.

### Pretest

A pre-test was conducted prior to the experiment to examine innate bias to low or high frequency tones. We compared between two groups with n=5 subjects each, in one the low frequency tones and in the other the high frequency tone was associated with the higher monetary gain in the discrimination condition (0.8 probability to gain money). The trials were divided into four periods, and the averaged selection probability was calculated for the tone that was associated with higher monetary gain. No bias was evident, and learning was similar to both options (Fig. 1D inset; Wilcoxon p>0.05 in each one of the periods).

### MR Scanning Protocol

MR data was collected using a 3 Tesla Tim Trio scanner (Siemens, Erlangen), with a 12-channel receiver head coil. Body coil with a peak B_1_ of 19 μT was used for transmission. Anatomical images were acquired using three-dimensional T_1_-weighted MPRAGE sequence (1×1×1 mm^3^ voxels, TR/TE/TI = 2300/2.98/990 ms, 176 slices, FA=9°, in-plane FOV=256×256 mm^2^, TA=4:44 min). The sagittal and coronal T_1_-weighted anatomical images enabled localization of a 40×25×10 mm^3^ ^1^H-MRS voxel on the midline of the dACC (Fig. 2A). In order to enhance magnetic field homogeneity, first and second order shim adjustments were performed using the default Siemens shimming tool, yielding water line widths of 6-7 Hz. The GABA and Glx ^1^H-MRS spectra were acquired using the spectral editing sequence Mescher-Garwood Point RESolved Spectroscopy (MEGA-PRESS)[8]. Each MEGA-PRESS scanning block (TR/TE=2000/68 ms, 2048 complex FID points, 2 kHz bandwidth, 16-step phase cycle) consisted of a metabolite scan (144 averages, TA=9:57 min) that was followed by a water reference scan (16 averages, TA=1:00 min). The individual coils’ spectra were phased and weighted by their signal to noise ratios using the reference coil sensitivity maps before combining their respective spectra.

### Procedure

Participants underwent initial training composed of four trials in front of a computer to ensure the task was correctly understood. Upon completion, they entered the magnet head first supine, while being visually monitored for awareness. A response box was handed to all subjects, and a projector was used for visual feedback while lying in the scanner. Anatomical images were acquired and enabled localization of the ^1^H-MRS voxel in the dACC. Shimming was carried out, followed by a ^1^H-MRS “rest” scan, during which the subjects were asked to focus on a fixation cross placed at the center of the screen. Following the rest scan, the participants began the experimental session in which, at the onset of each experimental condition block, a MEGA-PRESS metabolite scan was initiated, followed by a water reference scan. The responses of the subjects, along with the rewards they received, were recorded and labeled (Fig. 1B). The subjects were divided into two cohorts, each exposed to a different ordering of the experimental conditions (Fig. 1C): The first cohort (20 subjects) started with two blocks of the uncertainty condition, continued with one block of the null condition, and concluded with two blocks of the discrimination condition. The second cohort (11 subjects) started with one block of the null condition, followed by two blocks of the uncertainty condition and concluded with two blocks of the discrimination condition. Eighteen subjects (seven from the first cohort and all subjects from the second cohort) also performed an additional and final rest scan after concluding the task.

### Behavioral Analysis

The behavioral data was analyzed concentrating on the first block of each condition, using in-house MATLAB scripts. In order to examine the dynamics of the subjects’ choices over the course of each experimental condition’s block, we performed a moving average of the selection probability with a window width of 10 trials. Additionally, to assess behavioral performance per condition, we calculated the averaged selection probability of the high gain tone. To probe the learning rate in the discrimination condition, we calculated the mean overall selection probability, and used the first trial in which each subject reached this selection probability.

### 1H-MRS Processing and Quantification

MEGA-PRESS spectra were analyzed using in-house MATLAB scripts. Spectra were zero-filled 8-fold, apodized using an exponentially time decaying function with a linewidth of 3.2 Hz, Fourier transformed and phased. Difference spectra were generated by subtraction of the alternating ON and OFF spectra, which were subsequently aligned to the NAA methyl singlet at 2.01 ppm.

Metabolite quantification was achieved via peak integration with the integration limits 3.56 to 3.94 for Glx and 2.83 to 3.19 for GABA (Fig. 2B). The signals of GABA’s methylene group+ macromolecular contributions at 3.01 ppm [9, 53], and the Glx complex at 3.75 ppm are represented by:

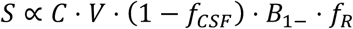

where C is the metabolite’s concentration, V the voxel volume, B_1_- the receiver coil sensitivity, f_CSF_ the fraction of CSF in the voxel, and f_R_ a factor accounting for T_1_ and T_2_ relaxation. Taking the ratio of each metabolite to the reference water signal (GABA/water, Glx/water) removed common factors such as B_1_- and V, and reduced signal variability. The remaining factors, such as *f_R_* and *f_CSF_*, as well as the macromolecular contamination, remained unaccounted for, but were assumed constant throughout the paradigm when interpreting our results. Since our analysis focused on intra-subject changes to metabolite levels during the learning task, these inter-subject sources of variability did not bias our conclusions. To further factor-out these constant elements we have also examined the standardized differences:

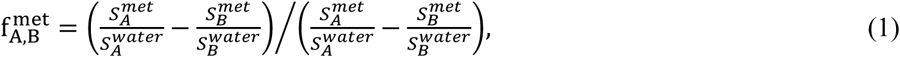

where *met*=Glx, GABA and A,B=null, discrimination, uncertainty or rest. The expression in Eq. (1) is independent of *f_CSF_*, *f_R_*, *B*_1–_ and any other factor assumed unchanged between conditions. This expression, when standardized to rest, served as standardization to baseline activity, and when was conducted between two conditions was reflecting a direct comparison between conditions of the task.

### Validating robustness of integration limits

In order to validate the robustness of choosing the integral limits for each metabolite, we further carried an optimization process, varying the integration limits, from 3.01±0.35 ppm for GABA and 3.755±0.35 ppm for Glx, and then doing multiple comparisons to choose the best ‘classifier’. In accordance to Eq. (1), the receiver operating characteristic (ROC) curve of each classifier was calculated three times, testing for a difference between two conditions of the task in each time (Fig. 2C inset). The areas under the curve (AUCs) of all the tested classifier were compared (Fig. 2C) and the classifiers with the highest AUC were selected in each comparison type. Subsequently, for each metabolite, the averages of the highest AUCs were set as the chosen integration limits: 3.56 to 3.94 for Glx and 2.83 to 3.19 for GABA. As a control, we performed this process on two sets of random noises. The AUCs for all the tested pairs were close to 0.5 (in gray line, Fig. 2C), implicating that no significant data could be found in case of noise, and that the integration limits that showed significance for the real data, indeed included important information. We note that the process resulted in integration limits that are highly similar to the ones chosen by an experienced observer and/or in similar studies that take such limits a-priori.

### Statistical tests

Changes to within-subject behavioral and metabolic measures were compared using two- or one-way repeated measure analyses. Tucky-Kramer tests were conducted for post-hoc paired comparisons. One sample t-tests was conducted in order to assess normalized metabolic levels significance, while paired sample t-tests was conducted for temporal metabolic drift analyzing. To analyze behavioral pretest data and in order to assess conditions ordering effect, we performed unpaired group comparisons using nonparametric Wilcoxon rank sum tests. Pearson’s correlation coefficient was conducted in order to test correlations between metabolites levels and behavioral performance. Unless otherwise stated, significance level was set to P<0.05.

